# Identification of a shared persistence program in triple-negative breast cancer across treatments and patients

**DOI:** 10.1101/2025.03.17.643634

**Authors:** Léa Baudre, Grégoire Jouault, Pacôme Prompsy, Melissa Saichi, Sarah Gastineau, Christophe Huret, Laura Sourd, Ahmed Dahmani, Elodie Montaudon, Florent Dingli, Damarys Loew, Elisabetta Marangoni, Justine Marsolier, Céline Vallot

## Abstract

Acquisition of resistance to anti-cancer therapies is a multistep process, which initiates with the survival of drug persister cells. Understanding the mechanisms driving the emergence of persister cells remains challenging, primarily because of their limited accessibility in patients. Here, using mouse models to isolate persister cells from patient tumors, we determine the identity features of persister cells from eight patients with triple-negative breast cancer (TNBC). Combining over 80 transcriptome studies, we reveal hallmarks of the persister state across patient models and treatment modalities: high expression of basal keratins together with activation of a stress response and inflammation pathways. Patient-derived persister cells are transcriptionally plastic and return to a common treatment-naïve like state upon relapse, regardless of the treatment they have been exposed to. Leveraging gene regulatory networks, we identify AP-1, NFKB and IRF/STAT as the key drivers of this hallmark persister state. As a proof of concept, we show that FOSL1 - an AP-1 member - is sufficient to drive cells to the persister state by binding enhancers and reprogramming the transcriptome of cancer cells. On the contrary, cancer cells without FOSL1 have a decreased ability to reach the persister state. By defining hallmarks of drug persistence to multiple therapies of the standard of care, our study provides a resource to design novel combination therapeutic strategies to limit resistance.

## Introduction

Acquiring a treatment-resistant cancer phenotype is a multi-step process^1^. Following exposure to an anti-cancer therapy, cancer cells can initially tolerate the treatment but then will undergo massive cell death, with only a few so-called persister cancer cells surviving. Initially non-cycling, a fraction of these persister cells will ultimately start cycling again^2^, thereby constituting a pool from which resistant cancer cells can emerge and actively grow under treatment^3,4^. Fully fledged resistance is driven by the acquisition of genetic alterations, with or without non-genetic alterations^5,6^. In contrast, persistence to cancer therapy is driven by non-genetic mechanisms, such as transcriptomic reprogramming^7^, epigenomic remodeling^8,9^, or translation deficiencies^10^; correspondingly, persistence was shown to be reversible *in vitro^2,8,11,12^*and *in vivo^13^*. In particular, in TNBC, persistence to therapy is driven by transcriptional plasticity^13^, metabolism reprogramming^14^, or alterations in DNA methylation or histone modifications^9,15,16^.

Numerous studies suggest that the switch from a treatment-naïve state to a persister state is the result of active reprogramming of a fraction of primed cancer cells, rather than the selection of a single clone. Lineage tracing studies show that the persister state is multi-clonal, and that a variety of treatment-naïve cancer cells can reach the persister state^9,13,17–19^. Following treatment exposure, a fraction of predisposed cells can reach the persister state^9,20^. However, as this pool of cells is constantly evolving, a potentially wider range of cancer cells can reach the persister state over a longer time^17,20^. Thus, while preventing this process holds promise as a therapeutic strategy for preventing the subsequent acquisition of drug resistance, we first require a detailed understanding of the mechanisms that drive the reprogramming of treatment-naïve cancer cells to persister cancer cells.

Transition to the persister state is known to be regulated by several chromatin modifiers, and in particular histone demethylase enzymes KDM5A/B^11,21–23^ and KDM6A/B^8,9^. However, these enzymes are hard to target with high specificity, limiting their use as a therapeutic option in humans. Other drivers of the switch from treatment-naïve to persister include receptors^7,24^, metabolic enzymes^25^, and transcription factors^7,26,27^. Targeting these drivers is currently under intense investigation as an option to enhance response to initial treatment and limit recurrence. Nonetheless, the range of patients who could benefit from these therapeutic options remains unknown. Indeed, the majority of studies have focused on in-depth understanding of mechanisms of drug persistence in *in vitro* model systems or in a few patient-derived models.

Characterizing the persister state in individual patients remains challenging. In the standard of care, biopsies are often performed pre-treatment (at diagnosis) or post–neoadjuvant treatment (after surgery). On-treatment biopsies, which could catch persister cells in the making, are rarely performed. Patient-derived xenograft (PDX) models represent a powerful strategy for isolating persister cells from whole tumors: tumor cells can be isolated during treatment as required, and their response to multiple treatments can be determined in parallel. To date, however, the lengthy process of generating PDXs has only been used to profile persister cells in three different patients tumors in parallel, in solid cancers: melanoma in response to BRAF-MEK^7^, TNBC in response to anthracyclin^13^ or capecitabine^9^, and colon cancer in response to Irinotecan^19^.

Here, we analyze the drug persister state across patients and therapies, based on persister cells derived from multiple patient samples at diverse steps of standard-of-care (SOC) treatment as well as with distinct treatment regimens. We find that no matter the cancer treatment, cancer cells in individual patient models reach a common persister state that is reversible and can be reset to the initial untreated state. We further identify a recurrent persister expression program across patients, potentially driven by a set of transcription factors, namely members of AP-1, IRF/STAT, and NF-kB complexes. We showcase the key role of these transcription factors in driving persistence with a detailed study of AP-1. We show that the presence of the AP-1–activating transcription factor FOSL1 is sufficient for the emergence of persister cells *in vitro* and that it drives the reprogramming of the expression program of cancer cells.

## Results

### Tumor cells from an individual patient consistently reach a common persister state that resets upon relapse

To isolate persister cells from patient samples, we first engrafted tumors on nude mice, and subsequently treated them with standard of care anti-cancer therapies. We retrieved persister cells by pooling the fat pads from mice once they showed pathological responses, as previously described (**Fig. 1A**)^9^. We leveraged this method to isolate patient-derived persister cells from eight TNBC PDX tumors (**Fig. S1A–C**), established either from treatment-naïve or residual tumors after patients had been exposed to several cycles of chemotherapy. We then treated mice with anthracyclines (AC) or platinum drugs, which are typically used in the neoadjuvant setting, or with capecitabine, the oral prodrug of 5-fluorouracil (5-FU), used in the adjuvant setting (**Fig. S1A**). For two PDXs (HBCx-33 and HBCx-14), we treated the mice independently with up to three chemotherapies (**Fig. 1B and Fig. S1B**). From a total of 127 mice, we performed transcriptomic analyses of *n* = 23 untreated tumors, *n* = 42 persister populations, and *n* = 22 relapsed tumors, using a combination of microarray, bulk RNA-seq, and single-cell (sc) RNA-seq to characterize the transcriptional program activated upon therapeutic stress *in vivo* (**Table 1**).

**Fig. 1.**
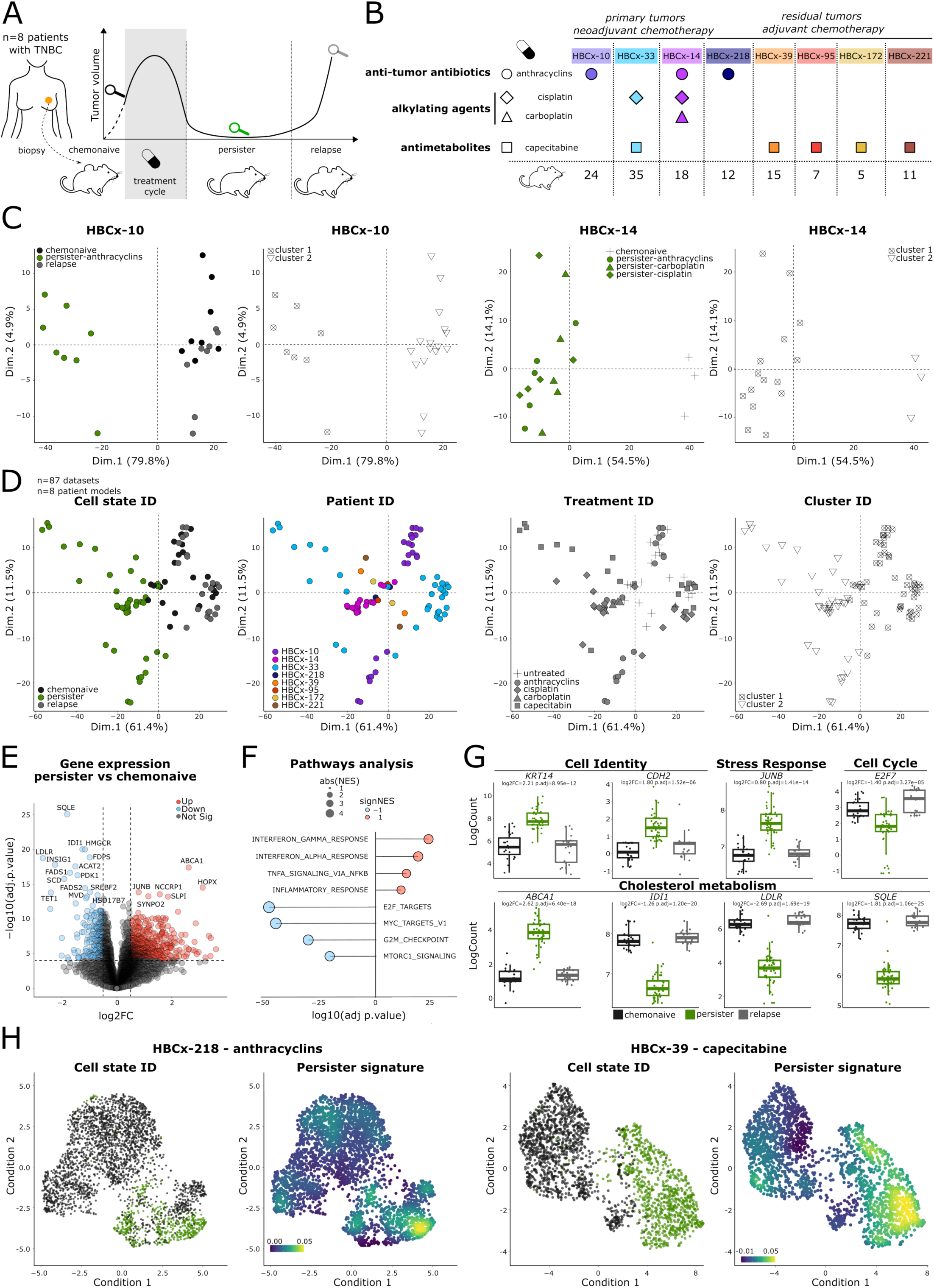
Identification of a recurrent and reversible persister state across TNBC patient models. **A.** Scheme of the PDX generation and treatment. **B.** Table of the models and chemotherapy treatments. The total numbers of mice analyzed in the study are indicated. **C.** PCA of the RNA dataset where samples are colored according to cell state ID or cluster ID, obtained for HBCx-10 (left) and HBCx-14 (right). **D.** PCA of the RNA dataset where all PDX samples are colored according to cell state ID, PDX model/Patient ID, treatment ID or cluster ID. **E.** Volcano plot representation of the gene expression differential analysis between chemo-naïve and persister cells in all PDX samples. **F.** Hallmark pathways activated (red) or downregulated (blue) in persister cells compared to chemo-naïve populations in all PDX samples. **G.** Boxplot representation of the expression of the indicated genes in chemo-naïve, persister and relapsed cells in all PDX models. Log2FC between chemo-naïve and persister cells and adj. *P* are indicated. **H.** UMAP of the scRNA-seq dataset obtained for HBCx-218 (left) and HBCx-39 (right) PDX models. Cells are colored according to the cell state ID or to the identified common persister signature.

We first analyzed the variability of the persister state in individual patient models, to determine whether a patient-specific tumor always gives rise to the same pool of persister cells. For this, we tested multiple mice (*n* = 16) engrafted with the same tumor of origin (HBCx-10) (**Fig. 1C**) after treatment with an AC chemotherapy, which is a combination of doxorubicin (a topoisomerase II inhibitor) and cyclophosphamide (an alkylating agent). After performing an unsupervised analysis of transcriptomes from all mice, principal component analysis (PCA), and hierarchical clustering, we observed a striking similarity between the eight persister populations derived from different mice: they all shared common transcriptional features that account for most of the variations observed in the collected transcriptomes (PC1 corresponds to 79.8% variance within all mice), and they all clustered together away from untreated tumors (**Fig. 1C**). Critically, the transcriptome of these persister populations always reset back to that of initial treatment-naïve cells upon relapse, as all relapsed tumors clustered with untreated tumors (**Fig. 1C**).

We further tested the observation that the persister populations are similar and reset upon relapse for another PDX (HBCx-33). Here, we observed that 11/14 persister populations clustered together and away from the group of untreated tumors (PC1 corresponds to 78.5% variance). Likewise, they reset to initial state upon relapse (*n* = 14 relapsed) (**Fig. S1D**). Mice were treated with two different chemotherapies: the alkylating agent cisplatin, and the anti-metabolite, capecitabine. Notably, transcriptomes from persister populations generated with either treatment were hardly distinguishable. To further investigate whether the persister state is treatment independent, we used a third model treated with three different chemotherapies: cisplatin, carboplatin (also an alkylating agent), and an AC (HBCx-14) (**Fig. 1C**). We found that all persister cell populations clustered together irrespective of the treatment. Altogether, we demonstrated *in vivo* that patient-derived cells recurrently reach a persister state that is common across therapies and resets to the same initial state upon relapse.

### Identification of a recurrent KRT14-positive persister state common across TNBC patient models and treatments

Next, we aimed to identify potential recurrent features of the persister state across patients and treatment regimens in TNBC. When combining datasets from all persister and control tumors (*n* = 87 datasets), we observed that persister states were first and foremost patient- or technology-specific (**Fig. S1E**). Thus, to search for recurrent features of the persister state across patient models, we next integrated datasets taking into account the patient and technology of origin^28^ (**Methods**). We showed that the main remaining source of variation among the 87 transcriptomes is the cell state, whether taken from persister, untreated, or relapsed cells (PC1 corresponds to 61.4% variance; **Fig. 1D**). Persister cell populations clustered together (39/42) away from untreated or relapsed cells, revealing the existence of a common and transient persister state across the different tumors and chemotherapy treatments (**Fig. 1D and Fig. S1F**). This recurrent consensus persister state is characterized by the activation of 836 genes (**Fig. 1E-G and Table 2**) with a log_2_ expression fold-change (herein, log2FC) over 0.5 and adjusted *P*-value (herein, adj. *P*) < 1e-4, which we defined as the consensus persister signature. Compared to untreated cells, persister cells displayed higher expression levels of basal genes (*KRT14, KRT6A*) and mesenchymal genes (*CDH2*), suggesting they are in a mixed epithelial and mesenchymal state (**Fig. 1G and Table 2**). These observations confirmed our initial findings on *n* = 3 patient models^9^ and suggest that these genes are hallmark markers of the persister state in TNBC independently of the chemotherapy regimen.

Persister cell populations all activated the interferon and inflammatory responses (*IRF/STAT*), and stress response (*JUNB*), while blocking the cell cycle (**Fig. 1E–G**). In addition, across patient models, persister cells showed a common drastic switch in cholesterol metabolism, with a combined shutdown of lipogenesis (*IDI1, SQLE*) and cholesterol import (*LDLR*) and a high expression of cholesterol efflux pump (*ABCA1*) (**Fig. 1G**). A major fraction of target genes of the SREBP2 transcription factor, which is responsible for the intracellular production of cholesterol, were repressed in persister cell populations across patient models; these repressed genes included *IDI1, LDLR, SQLE, FDPS,* and *HMGCR* (**Fig. 1E-G and Table 2**). Altogether, we show that in TNBC, across treatment regimens, patient-derived persister cells share a common basal and mesenchymal identity, with stress-associated features and an alternative cholesterol metabolism.

We identified high KRT14 expression as one of the hallmarks of the persister state in TNBC. Of note, KRT14 is used clinically as a prognostic marker to characterize breast cancers (in conjunction with KRT5/6, estrogen (ER), progesterone (PR), human epidermal growth factor receptor-2 (HER2) and Ki-67). KRT14 and/or KRT5/6 define the basal-like phenotype within TNBC subtypes that do not express ER, PR, or HER2^29^ and are associated with a poor outcome^30^. Interrogating the PanCancer Breast cancer atlas comprising chemo-naïve tumors^31^, we observed that basal-like tumors express KRT14 levels close to that of the normal adjacent tissue, which contains basal cells. In addition, about a fourth of these tumors (24%) express KRT14 at levels that are higher than that of the adjacent normal tissue (**Fig. S1G**). These tumors are associated with lower disease-free survival rates than other basal-like tumors (*P* = 3.10e-2) (**Fig. S1H**). These KRT14-high tumors could contain a subset of KRT14-positive persister-like or pre-persister cells prior to treatment.

Focusing on two patient-derived models (PDX HBCx-218 and -39), we next quantified the intra-tumor heterogeneity of the identified consensus persister state using scRNA-seq. For both PDX models, we detected activation of our newly-defined consensus persister gene signature in persister cells, whereby it was high in most persister cells for HBCx-39 and high in only a fraction of the persister cells in HBCx-218 (**Fig. 1H**). This suggests that part of the persister cell populations activate patient model-specific genes that are not part of the consensus persister gene signature. Further, we detected low activation of the persister gene signature in untreated tumors, suggesting that at least part of the genes are already expressed prior to treatment, as we observed for KRT14 in human samples.

### Transient activation of AP-1, NF-kB, and STAT/IRF transcription factors in persister state

We next inferred regulon activity to predict the regulators of the persister expression signature (**Fig. 2A**). Using the decoupleR framework^32^ (**Fig. 2A**), we show that the top 10 regulons explaining the consensus persister expression signature are associated with NF-kB (*NFKB1, RELA*), IRF/STAT (*STAT1/IRF1*), and AP-1 (*JUN*). In addition, a series of individual transcription factors (*ETS1*, *CNNTB1,* and *SPI1*) were also predicted as potential regulators of the consensus expression signature. The predicted regulons are transiently activated in persister cells and are then shut down in cells at relapse (**Fig. 2B**). These predictions match the pathways that we have found to be characteristic of the consensus persister state. Transcription factors of the IRF and STAT families are indeed orchestrators of the interferon response^33,34^, while the AP-1 and NF-kB complexes are major actors of stress-response^35–37^. Differences in regulon activity could result from changes in the expression or activity of transcription factors. Analyzing our transcriptome datasets, we found that only a fraction of these transcription factors are overexpressed in persister cells (**Fig. 2C, Fig. S2A-C, and Table 2**). Among the AP-1 family, *JUNB* is the most recurrently overexpressed member across patient models and treatments (log2FC = 0.80, adj. *P* = 1.41e-14), while *RELB* is the most overexpressed genes within the NF-kB family (log2FC = 0.50, adj. *P* = 2.54e-5), and *IRF7* (log2FC =1.49; adj. *P* = 2.12e-9) and *STAT1* (log2 FC = 0.84; adj. *P* = 2.10e-6), within the IRF/STAT families. These changes in expression could explain in part the regulon activities observed in the persister state. Using the TCGA PanCancer atlas, we analyzed the patterns of expression of these transcription factors prior to treatment. As a follow-up of our previous observation on *KRT14*-high chemo-naïve tumors, we observed that *FOSL1* and (to a lesser extent) *JUNB* are the two transcription factors that most significantly correlated with *KRT14* (correlation score= 0.51 and 0.39; *P* = 3.20e-12 and 1.70e-7, respectively; **Fig. 2D**). This observation suggests that *KRT14*/AP-1–high tumors exist prior to treatment.

**Fig. 2.**
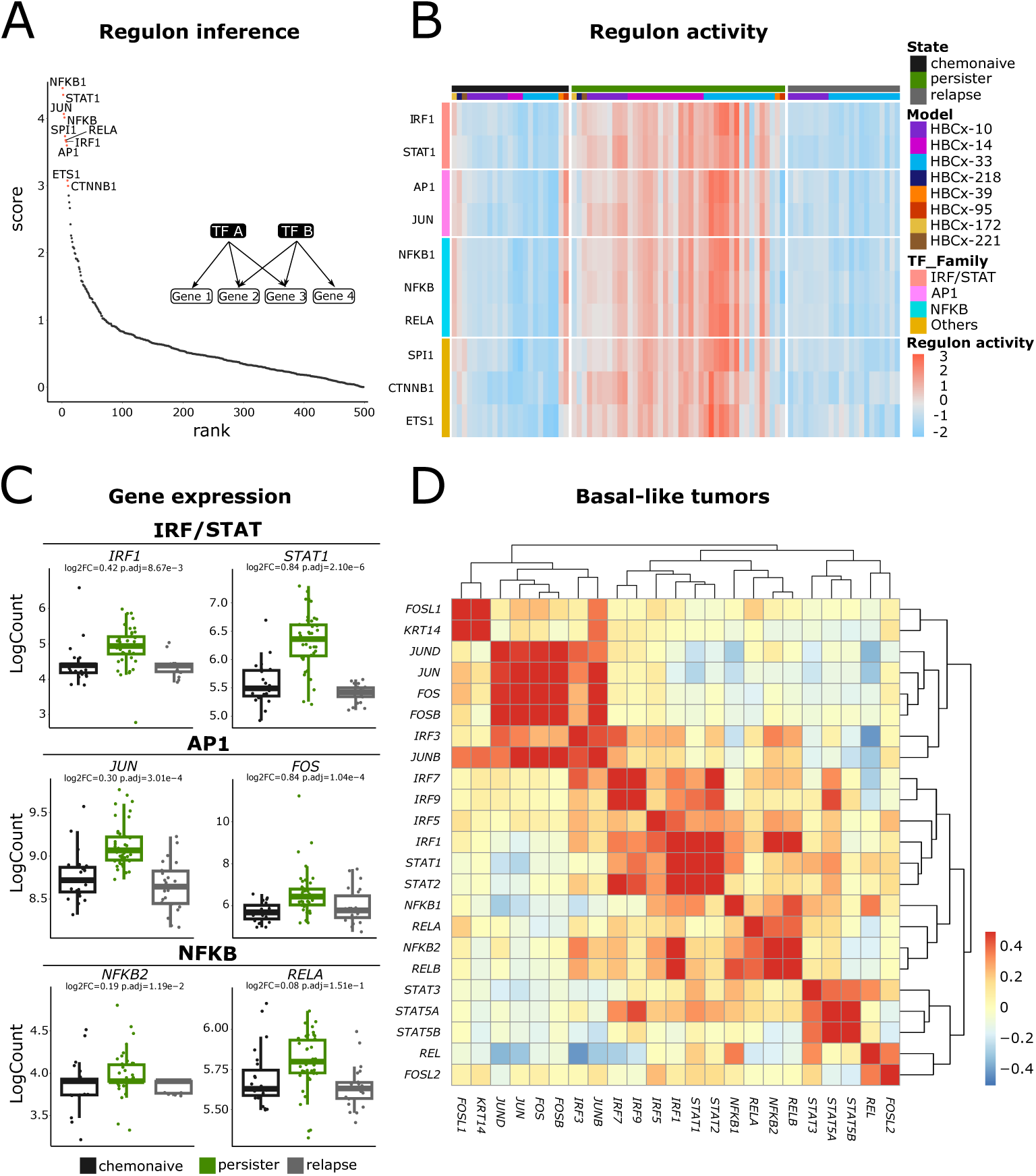
Recurrent transcription factors regulate the drug persister expression program in TNBC. **A.** Scoring of regulon activity inference on the common persister program. Top 10 scoring transcription factors are highlighted in red. **B.** Heatmap representation of the regulon activity inference of the indicated predicted transcription factors in each PDX model. **C.** Boxplot representation of the gene expression of the indicated transcription factors in chemo-naïve, persister, and relapsed cells in all PDX models. Log2FC between chemo-naïve and persister cells and adj. P are indicated. **D.** Heatmap representation of the pairwise correlation between the normalized values of the persister gene *KRT14* and the predicted transcription factors in human basal like-tumors form the PanCancer Breast cancer atlas.

### Remodeling of the H3K27ac epigenome upon drug insult

To study the mode of action of the potential TF drivers we identified, we first tested *in vitro,* whether TNBC cell lines could recapitulate at least part of the consensus persister state and the transcriptional regulons we observed in PDX samples (**Fig. 3A**). We performed scRNA-seq on three different TNBC cell lines (MDAMB468, HCC38, and BT20) treated with chemotherapy (5-FU) and identified a set of genes common to all persister cell populations (**Fig. S3A and B**). To characterize the regulatory landscape of the persister state, we used inference of regulon activity and observed that part of the top 10 regulons potentially driving a persister program *in vitro* belong to NF-kB (*RELA*) and AP-1 (*JUN*) regulons (**Fig. 3B**). This regulatory network recapitulates in part what we observed in patient-derived persister cells obtained from our TNBC PDX models.

**Fig. 3.**
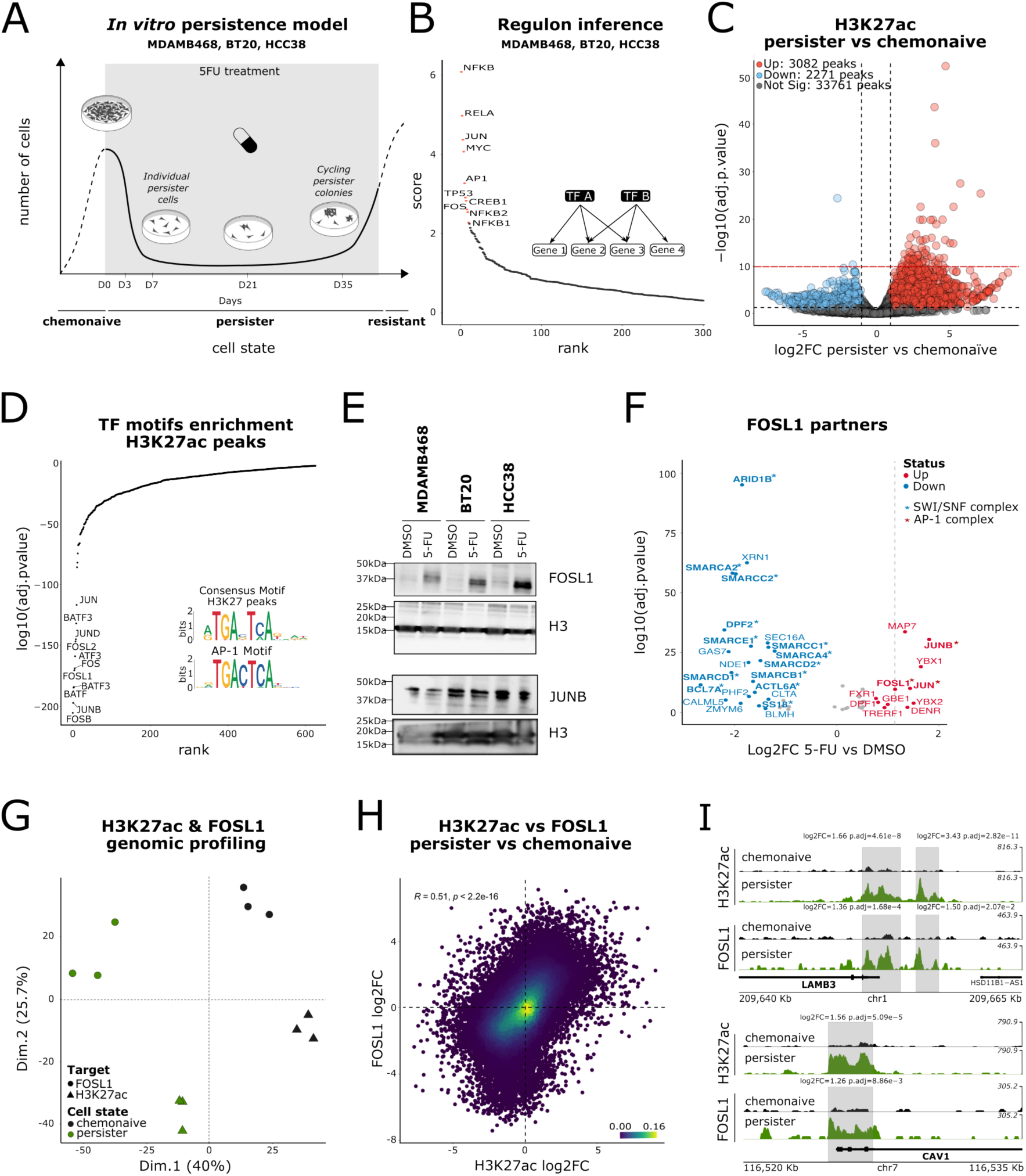
Epigenomic remodeling upon treatment is dedicated to AP-1 sites. **A.** Representative scheme of the evolution of TNBC cell lines under 5-FU treatment. Timepoints and associated phenotypes observed at the microscope are represented. **B.** Scoring of regulon activity inference on the persister program of 3 TNBC cell lines, MDAMB468, BT20 and HCC38. Top 10 scoring transcription factors are highlighted in red. **C.** Volcano plot representation of the differentially enriched H3K27ac peaks in persister cells treated 35 days with 5-FU compared to chemo-naïve cells in MDAMB468 cells. Red horizontal line represents an adj. *P* < 1e-10. **D.** Representation of the transcription factors binding motifs enriched within H3K27ac gained peaks in persister cells (35 days of treatment) compared to chemo-naïve MDAMB468 cells. Identified consensus motif in H3K27ac peaks and genomic binding motif of AP-1 are represented. **E.** Representative images of the protein expression of the two AP-1 canonical members: FOSL1 and JUNB in the indicated cell lines treated for 21 days with 5-FU. Histone H3 is used as a loading control (*n* = 3 biological replicates). **F.** Volcano plot representation of the differentially enriched peptides in FOSL1 immunoprecipitation between 5FU and DMSO treatment for 7 days (MDAMB468 cells, n=5 replicates, n=84 proteins). The vertical dashed line corresponds to the log2FC for the FOSL1 protein. **G.** PCA representation of H3K27ac distribution or FOSL1 genomic binding profiles in MDAMB468 chemo-naïve and persister cells treated for 35 days with 5-FU. **H.** Scatterplot representation of FOSL1 vs H3K27ac log2FC upon 5-FU treatment in MDAMB468 cells treated for 35 days. Log2FC corresponds to ratios of H3K27ac enrichment persister between persister and chemo-naïve cells. Pearson’s correlation scores and associated *P*-value are indicated. **I.** H3K27ac and FOSL1 profiles over LAMB3 and CAV1 loci in MDAMB468 chemo-naïve and persister cells (35 days of treatment with 5-FU). Log2FC enrichments between chemo-naïve and persister cells and corresponding adj. *P* are indicated.

To further understand the epigenomic mechanisms driving transcriptional changes, we used H3K27ac profiling to map active enhancers in chemo-naïve or persister cells from the MDAMB468 cell line (**Fig. 3C and Fig. S3C-E**). The persister state was characterized by a rewiring of the H3K27ac landscape, with the most significant changes corresponding to localized gains of H3K27ac (*n* = 106 enriched peaks vs *n* = 4 depleted peaks, with adj. *P* < 1e-10). These changes in chromatin are associated to the activation of expression of the nearest genes (**Fig. S3D**) in persister cells. To understand which transcription factors could be associated with the chromatin remodeling at these peaks, we performed motif enrichment on peaks enriched for H3K27ac after treatment (**Fig. 3D**). We found that the AP-1 related motifs were by far the most significantly enriched in these persister cells. Thus, we focused on the AP-1 regulon for the rest of the study.

### The FOSL1 protein is overexpressed upon treatment, switches partners and accumulates at enhancers together with H3K27ac

We next studied the expression level of the AP-1 members *in vitro* upon chemotherapy treatment. At the protein level, we detected significant changes in the expression of the core AP-1 members JUN and FOSL1 (**Fig. 3E and Fig. S3F**). Specifically, both are upregulated in all three cell lines upon 21 days of treatment, while JUNB, JUND, and FOSB are expressed at similar levels in chemo-naïve cells and persister cells (note that no suitable antibodies were available for the other AP-1 members). As FOSL1 displayed the highest fold-change in expression (average FC = 6.7 in protein expression in all three cell lines), we focused our efforts on this AP-1 member. We showed that FOSL1 is highly expressed starting at day 7, with a slight activation detected already at day 3 (**Fig. S3G**). Notably, after chemotherapy exposure, day 7 corresponds to the time point at which a large fraction of the cell population has died, with only persisters left (**Fig. S3G**). Altogether, these results suggest that FOSL1 is highly expressed in persister cells at their emergence.

In addition to the activation of FOSL1 expression we investigated the evolution of the repertoire of FOSL1 partners upon 5-FU with mass-spectrometry (**Fig. 3F, Fig. S3H**). Consistent with FOSL1 overexpression upon treatment, we observed a significant increase of FOSL1 peptides after 5-FU. Ratio between FOSL1 peptides and several partners were increased upon treatment: we detected higher levels of YBX1-2 and JUNB and JUN peptides compared to FOSL1, indicating that interaction with these proteins is significantly enriched independently of the gain in FOSL1 expression. On the contrary, interaction of FOSL1 with several members of the SWI/SNF family - expected to form a canonical BAF complex - are lost upon treatment. Altogether, these results indicate that FOSL1 could be switching partners upon treatment.

Next, to determine whether AP-1 factors, particularly FOSL1, are involved in the regulation of the persister expression program, we first used *in situ* tagmentation (CUT&Tag) to profile FOSL1 binding sites in persister cells, and compared FOSL1 and H3K27ac profiles. Applying PCA on a common consensus peak annotation, we demonstrated that the H3K27ac and FOSL1 genomic binding patterns are extremely similar, grouping together on the first principal component of maximal variation (**Fig. 3G**). In particular, changes at genomic peaks in H3K27ac and FOSL1 enrichments upon treatment were significantly correlated (**Fig. 3H**). Chromatin remodeling upon treatment was characterized by the simultaneous enrichment of FOSL1 and H3K27ac on persister genes in persister cells as compared to those in chemo-naïve cells (**Fig. 3I and Fig. S3I**). Indeed, we found that 26% of all persister genes in MDAMB468 cells gained both FOSL1 and H3K27ac upon 5-FU treatment, suggesting that this AP-1 member is a key regulator of the persister expression program.

### FOSL1 is sufficient to reach the persister state and FOSL1 KO cells hardly reach the persister state

To test whether FOSL1 is sufficient to reach the persister state *in vitro*, we generated gain-of-function (GOF) models that constitutively overexpress FOSL1. Specifically, MDAMB468 cells were infected with a tagged *FOSL1* ORF (‘HA-FOSL1’) or with a CRISPR-Cas9 VPR system and two different gRNAs targeting *FOSL1* (‘gRNA1’ and ‘gRNA2’ cell populations; **Fig. 4A-C, Fig. S4A**), or respective control constructs (**Methods**). We selected (via the fluorescent reporter genes) for transduced cell populations, rather than individual clones, to retain maximum heterogeneity of the population. Indeed, depending on the site of integration of the cassette (tagged FOSL1 ORF or gRNAs targeting FOSL1), cells can express variable levels of FOSL1 protein (**Fig. 4C**). We next measured global expression levels of the FOSL1 protein in the various cell populations. HA-FOSL1 cells express high levels of *FOSL1* as compared to gRNA1 and gRNA2 cell populations, while cells with the CRISPR-Cas9 activating system, which activates the endogenous *FOSL1* promoter, have expression levels closer to that of 5-FU–exposed cells (**Fig. 4B-C, Fig. S4A**).

**Fig. 4.**
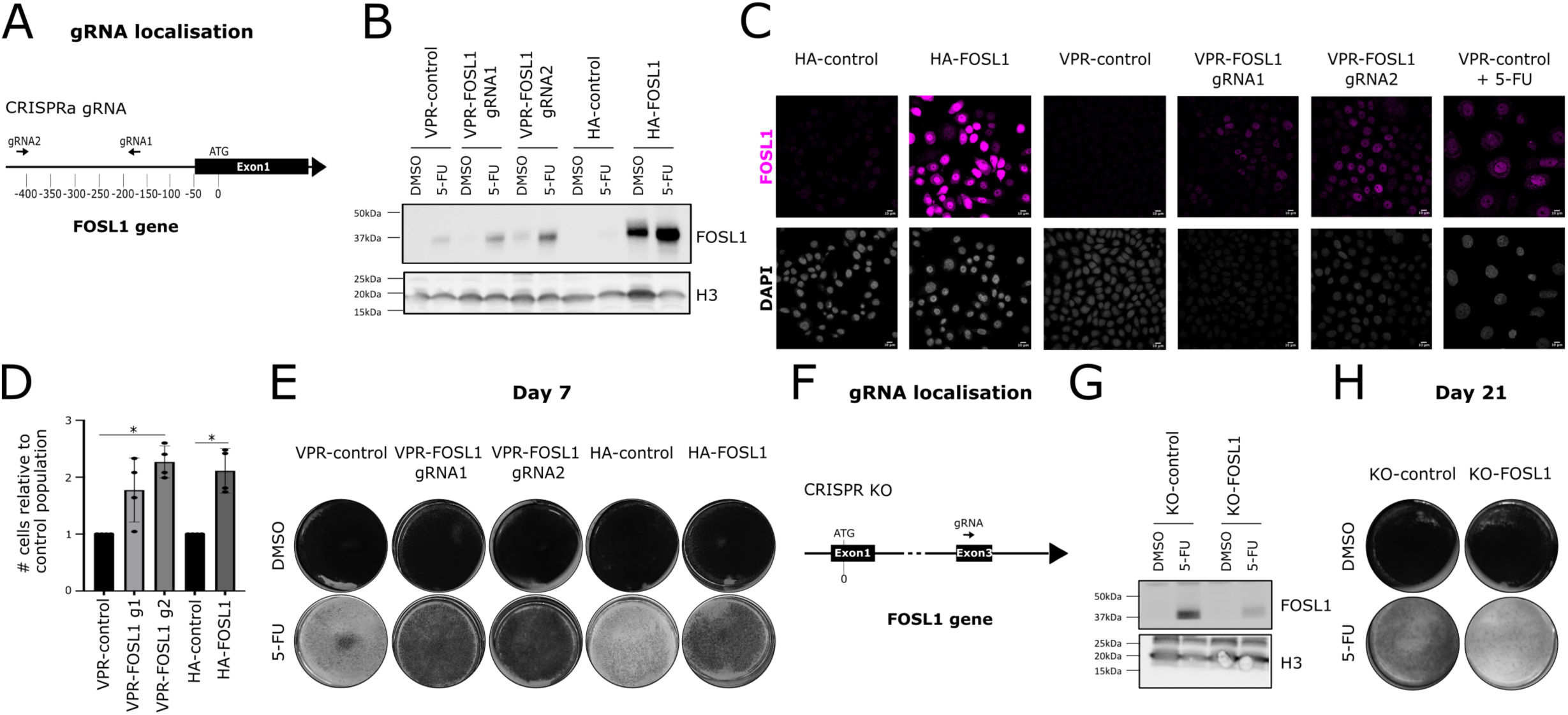
FOSL1 is sufficient to reach the persister state and FOSL1 KO cells cannot reach the persister state. **A.** Scheme representation of the genomic location of CRISPR VPR gRNAs at the FOSL1 locus. **B.** Protein expression of FOSL1 in the indicated MDAMB468 GOF cell lines treated 7 days with 5-FU Histone H3 is used as a loading control. Representative images are shown (*n* = 3 biological replicates). **C.** FOSL1 protein localization in the indicated MDAMB468 GOF cell lines treated or not with 5-FU for 7 days. Representative images are shown. **D.** Histogram representing the number of MDAMB468 cells after 7 days of treatment with 5-FU relative to DMSO (*n* = 3; mean, ANOVA, Kruskal-Wallis test; \**P* = 0.05). **E.** Colony forming assays for 5-FU–treated FOSL1 MDAMB468 GOF cells after 7 days of treatment with 5-FU. Representative images are shown (*n* = 3 biological replicates). **F.** Scheme representation of the genomic location of the CRISPR KO gRNA at the FOSL1 locus. **G.** Protein expression of FOSL1 in the indicated MDAMB468 KO cell lines treated 21 days with 5-FU Histone H3 is used as a loading control. Representative images are shown (*n* = 3 biological replicates). **H.** Colony forming assay for 5-FU–treated FOSL1 MDAMB468 KO cells after 21 days of treatment with 5-FU. Representative images are shown (*n* = 3 biological replicates).

We then evaluated the phenotypic consequences of *FOSL1* expression by comparing the ability of GOF cells to survive a 7-day 5-FU treatment. Notably, activation of *FOSL1* had no effect on any of the untreated cancer cell constructs, cells overexpressing FOSL1 proliferate similarly to control cells (**Fig. 4D-E and Fig. S4B**). In contrast, it significantly increased the survival of the 5-FU–treated cells as compared to the respective non-induced control populations (by 2.11 fold in HA-FOSL1 cells, and by 2.27 fold in gRNA2 cells; **Fig. 4D**). These results suggest that even a low expression of the FOSL1 protein is sufficient to confer the cells with the ability to survive chemotherapy, without stopping the cell cycle. In contrast, preventing the activation of *FOSL1* expression upon treatment using a CRISPR knockout (KO) strategy decreased the number of persister cells at day 21 (**Fig. 5F-H, Fig. S4C-D**). Thus, we conclude that FOSL1 is crucial in the transition from chemonaive to persister phenotype.

**Fig. 5.**
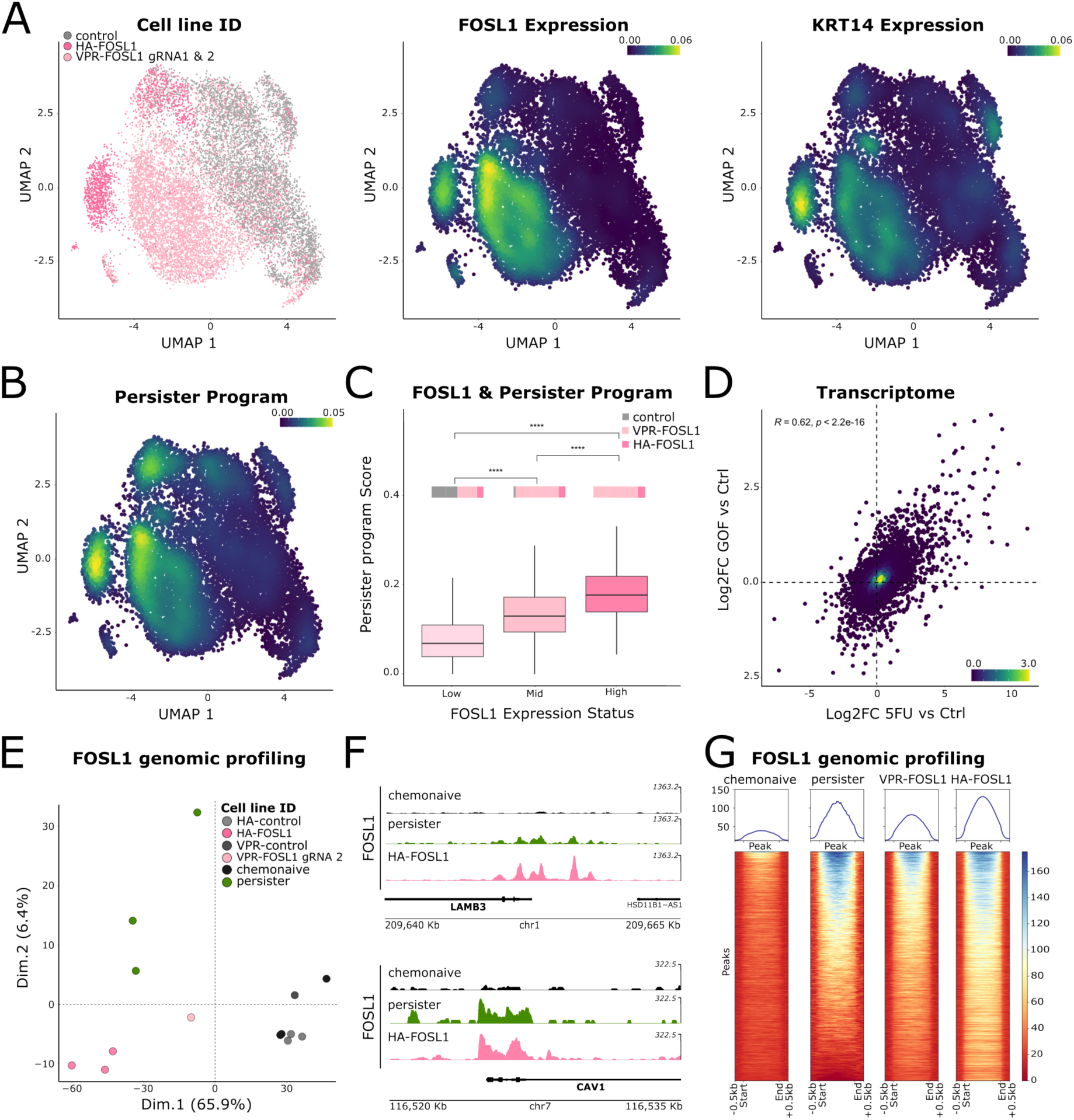
FOSL1 drives the activation the persister expression program in chemo-naïve cells independently of treatment. **A.** UMAP representing the MDAMB468 FOSL1 GOF scRNA-seq dataset, colored according to sample of origin (left), FOSL1 (middle), or KRT14 (right) gene expression. **B.** UMAP representation of scRNA-seq dataset generated from MDAMB468 FOSL1 GOF populations, where cells are colored according to the level of activation of the identified persister expression program. **C.** Boxplot representing the level of activation of the persister expression program compared to the FOSL1 expression status in MDAMB468 control and FOSL1 GOF cell lines. The repartition of the samples in the FOSL1 expression status categories are represented on the top of the graph. *P* value of t-test comparison between FOSL1 Expression Status groups is indicated on the top of the graph. **D.** Dotplot representing log2 expression fold-change induced by 5-FU treatment^9^ or FOSL1 overexpression in MDAMB468 cells compared to the corresponding control population. Pearson’s correlation scores and associated *P* are indicated. **E.** PCA representation of FOSL1 genomic binding profiles in MDAMB468 cells treated 35 days or not with 5-FU and control or FOSL1 GOF cells. **F.** FOSL1 profiles over LAMB3 and CAV1 loci in MDAMB468 chemo-naïve, persister (35 days) and HA-FOSL1 cells. **G.** Heatmap representation of signal in FOSL1 peaks in MDAMB468 chemo-naïve, 5-FU–treated persister cells (35 days), and FOSL1 GOF cells. FOSL1 peaks in persister cells were used as reference and ordered by decreasing signal. The average levels of FOSL1 signal for the indicated samples are presented above.

### FOSL1 can drive the activation of the persister expression program in chemo-naïve cells independently of treatment

To investigate how FOSL1 induces the persister state, we analyzed the transcriptome and epigenome of cells with an exogenous activation of *FOSL1* expression in the absence of chemotherapy. We first analyzed the effect of different levels of *FOSL1* expression on the expression programs of cancer cells using multiplex scRNA-seq analysis. As expected, VPR-FOSL1 (gRNA1 & gRNA2) and HA-FOSL1 cell populations showed heterogeneous *FOSL1* gene expression levels (**Fig. 5A**). HA-FOSL1 cells (with highest protein expression of *FOSL1*) fully resembled 5-FU–treated persister cells (**Fig. S5A**), with the maximal expression of the *KRT14* gene (**Fig. 5A**), which is characteristic of the persister state *in vivo* and *in vitro^9^*. Increasing *FOSL1* expression levels led to a gradual shift of the expression program toward the persister state (**Fig. 5B-C and Fig. S5B**): *FOSL1*-low cells activated low levels of the persister expression program as compared to mid- or high-level expressing cells. *FOSL1*-low cells correspond to control cell populations, in which *FOSL1* is lowly expressed in absence of chemotherapy, or to a fraction of VPR-FOSL1 or HA-FOSL1 cells in which *FOSL1* mRNA is not detected. Altogether, expression changes induced by *FOSL1* activation are significantly similar to changes induced by 5-FU chemotherapy (**Fig. 5D**), suggesting that *FOSL1* activation alone can recapitulate the persister state without chemotherapy exposure.

In parallel, we investigated the genomic binding profile of FOSL1 in cells with forced *FOSL1* activation in the absence of chemotherapy. We showed that *FOSL1* activation recapitulated the FOSL1 binding pattern induced by chemotherapy exposure (**Fig. 5E-G**), showing that FOSL1 is able to reach its target genes in absence of therapeutic stress. Increasing levels of the FOSL1 protein led to a wider range of target sites (**Fig. 5G and Fig. S5C**). Notably, the H3K27ac landscapes of cells with activated *FOSL1* expression were similar to that of chemo-naïve cells (**Fig. S5D**), with no H3K27ac changes even at genes with induced expression (**Fig. S5E-F**). These results suggest that pre-existing levels of H3K27ac could be sufficient for FOSL1 to activate gene expression. Other histone modifications or chromatin modifiers could also facilitate gene activation by FOSL1.

## Discussion

By using eight PDXs from different patients with TNBC, which we treated with up to three different chemotherapies in parallel, we identified recurrent features of the persister state in TNBC. We observed that persistence to chemotherapy is a transient and reversible state, as has been previously described in other cancers^2,7,8,11,13^. We further show that persister cells derived from an individual patient model shared common features across different treatments and reverted back to a common initial state at relapse; thus, part of the persister expression program is patient-specific. Nonetheless, we identified general hallmarks of persister cells in TNBC shared across patient models irrespective of the treatment. These hallmarks are a high level of expression of basal keratins, a switch in cholesterol metabolism, and the activation of the interferon as well as of the stress responses, whereby the latter being in agreement with previous studies investigating early response to chemotherapy in breast and ovarian cancers *in vivo^38,39^*.

We identified KRT14 as a recurrent marker of persister cells *in vitro* and *in vivo* in TNBC. Whether KRT14 is a sole marker of basal-like identity or has a functional role in the persister phenotype remains to be investigated. So far, studies have demonstrated that KRT14 can mediate tumorigenesis and invasion in breast cancer^40^, ovarian cancer^41^, and bladder cancer^42^. Furthermore, a high KRT14 expression is closely associated with advanced tumor stage and poor prognosis in melanoma^43^ and ovarian cancer^41^ and may regulate the treatment response to cancer therapeutics in bladder cancer^44–46^. Here, we show that prior to treatment, some tumors expressed extremely high levels of KRT14, associated with the expression of the AP-1 members FOSL1 and JUNB. The patients from which these tumor cells were extracted showed a lower disease-free survival, suggesting that pre-existing persister-like features could be associated with poor response to neo-adjuvant chemotherapy and high recurrence rates.

Through epigenomic and transcriptomic analyses of persister cells, we identified the AP-1 complex as a key regulator of the persister state. We showed that FOSL1 is sufficient to launch the persister expression program, and that it appears necessary, in the model we tested, for cells to reach the persister state, indicative of a regulatory role in the switch to persister state. FOSL1 was previously shown to regulate the epithelial-to-mesenchymal (EMT) transition in aggressive breast cancer cells^47–49^ and to lead to shorter metastasis-free survival in patients with basal-like breast cancer^50^. More generally, several studies have highlighted the role of the AP-1 complexes in response to chemotherapy or to immuno- or targeted-therapies in cases of melanoma^20,51–53^, lung^54^ or ovarian cancers^55^. Here, we show that AP-1 might be a general regulator of the persister state in TNBC by characterizing the persister cells in response to multiple treatments.

We had previously shown that the activation of persister genes is accompanied by the loss of the Polycomb associated mark H3K27me3 at promoters^9^. Here we further show that H3K27ac is actually enriched at a set of these loci upon treatment, suggesting an enhancer rewiring upon the switch to the persister state. Transcription factors, including AP-1, might facilitate the chromatin remodeling observed at persister genes. Consistently with this hypothesis, AP-1 transcription factors have been recently described as pioneer factors capable of rewiring 3D enhancer landscapes together with the multimeric chromatin remodeler SWI/SNF complex^56,57^ or by cooperating with chromatin remodelers, such as CBP/p300^58^. Investigating additional histone marks and DNA methylation distribution dynamics in response to treatment will be instrumental to better understand the intimate link between chromatin remodeling and identified TFs activity in response to anticancer therapies.

We predict that other families of transcription factors could also regulate—together or not with AP-1—the recurrent persister program we identified, such as members of the IRF/STAT and NF-kB families, which are known to contribute to the therapeutic stress response in some cancers^39,59–63^. Further mechanistic investigations are needed to demonstrate whether transcription factors actually drive persistence in TNBC alone or in combination. Such cooperation of AP-1 with other transcription factors has interestingly been described for therapy resistance in prostate cancer and melanoma, with AP-1 factors driving phenotype switching together with the YAP/TAZ-TEAD complex^64,65^. The co-occupancy between AP-1 factors and these partner transcription factors on chromatin has also been experimentally demonstrated in breast cancer cells^66^, and loss-of-function of the partner transcription factor is sufficient to prevent AP-1-driven tumorigenesis *in vivo*. Therefore, inhibiting the interaction between transcription factors might offer additional opportunities to prevent its activity. Specific pharmacological compounds against transcription factors are not yet available given the absence of catalytic activities or binding pockets for small-molecule inhibitors^67,68^. New strategies like degraders-based technology are currently being developed to eliminate TFs in cancer cells^69,70^. They could represent an opportunity, in combination with SOC in TNBC, to prevent the emergence of persister cells and delay recurrence. SOC in TNBC has recently evolved with the inclusion of immunotherapies in neo-adjuvant treatments. We will need to understand, using immuno-competent models, which are the most relevant vulnerabilities of persister cells in this treatment scheme.

## Materials and Methods

### Experimental approaches

#### PDX models

Eight xenograft models were used that had been previously generated from eight different patients with TNBC, whereby HBCx-10, HBCx-14, and HBCx-33 were generated from primitive tumors, and HBCx-218, HBCx-39, HBCx-95, HBCx-172, and HBCx-221 were generated from residual tumors post–neoadjuvant chemotherapy. All models were established at the Curie Institute with informed consent from the patient. Female Swiss nude mice were purchased from Charles River Laboratories and maintained under specific pathogen-free conditions. Mouse care and housing were in accordance with institutional guidelines and the rules of the French Ethics Committee (project authorization no. 02163.02).

As indicated, mice were treated with various methods: capecitabine (Xeloda; Roche Laboratories) at a dose of 540 mg/kg, orally 5 d/week, for 6 to 14 weeks; adriamycine (Accord) at 2 mg/kg, intraperitoneal (i.p.) 1×/21 days; cyclophosphamide (Baxter) at 100 mg/kg, i.p. 1×/21 days; cisplatin (Mylan) at 6 mg/kg, i.p. 1×/21 days; or carboplatine (Accord) at 90 mg/kg, i.p. 1x/21 days. Relative tumor volumes (mm^3^) were measured as described previously. Before downstream analysis, cell suspensions were prepared as previously described^38,9^.

#### Cell lines, culture conditions, and drug treatments

MDAMB468 cells were cultured in DMEM (Gibco-BRL, ref. 11966025), supplemented with 10% heat-inactivated fetal calf serum (Gibco-BRL, ref. 10270-106). The HCC38 and BT20 cell lines were cultured in RPMI 1640 (Gibco-BRL, ref. 11875085), supplemented with 10% heat-inactivated fetal calf serum. All cell lines were cultured in a humidified 5% CO_2_ atmosphere at 37°C, and were shown to be mycoplasma negative. Cells were treated with 5 μM of 5-FU (Sigma, ref. F6627) for 7 to 21 days, as indicated.

#### Cloning

FOSL1 gRNAs were selected from the Weissman library^71^. Forward and reverse oligonucleotides were synthesized by Integrated DNA Technologies - IDT and cloned in pJR85-gRNA-BFP (Addgene, ref. 140095) using the BstXI and BlpI restriction enzymes.

FOSL1 cDNA was mutated by adding an HA tag after the ATG site and cloned by Genscript in pHIV-SFFV-MCS-IRES-GFP using BamHI and XhoI restriction enzymes.

#### Lentivirus packaging and cell transduction

Cas9WT-P2A-mCherry (Addgene, ref. 99154) and dCas9-VPR-P2A-mCherry (Addgene, ref. 154193) were used. gRNA lentivirus particles was produced by transfecting the plasmids pJR85-gRNA-BFP (Addgene, ref. 140095), psPAX2 (Addgene, ref. 12260), and pMD2.G (Addgene, ref. 12259) into HEK293T cells (ATCC® CRL-3216™) with PEIpro (Polyplus, ref. 101000017) and NaCl, following manufacturer instructions. Lentiviruses were titrated after infecting HEK293T cells with cascade dilutions and assessing the percentage of BFP fluorescence on a cytometer. MDAMB468 cells from ATCC at passage 11 were infected with lentivirus produced at a low multiplicity of infection (MOI 0.1). Ten days after transduction, cells were sorted for mCherry expression to select cells with Cas9WT or dCas9-VPR - mCherry expression. These cell lines were then transduced with FOSL1 gRNA lentivirus at MOI 0.3, and transduced BFP+ cells were isolated three weeks after transduction.

For tagged cDNA overexpression, lentivirus particles were produced by transiently transfecting the pHIV-SFFV-MCS-IRES-GFP or the HA-FOSL1-pHIV-SFFV-MCS-IRES-GFP into HEK293T cells using the calcium-phosphate transfection method. Lentiviral constructs were co-transfected with pMD2.G (Addgene, ref. 12259) and psPAX2 (Addgene, ref. 12260) (both kindly provided by Didier Trono). The culture media was collected 72 h after transfection of HEK293T cells. Lentiviral particles were concentrated by ultracentrifugation. Lentiviruses were titrated after infecting HEK293T cells with cascade dilutions and assessing the percentage of GFP fluorescence on a cytometer. MDAMB468 cells from ATCC at passage 2 were infected with lentivirus produced at a low multiplicity of infection (MOI 0.5). Ten days after transduction, cells were sorted for GFP expression to select cells with GFP expression or HA-FOSL1-GFP expression.

In all cell lines, expression of FOSL1 was validated at both the protein and mRNA level.

#### Western blot

Cells lysates were prepared as previously described^9^. Antibodies anti-H3K27ac (dilution 1:1000; Cell Signaling, ref. 9733), FOSL1 (Dilution: 1:500, Santa Cruz, ref. sc-28310), JUN (dilution 1:100; Abcam, ref. ab31419), FOS (Dilution: 1:500, Cell Signaling, ref. 2250), FOSB (Dilution: 1:500, Cell Signaling, ref. 2251), JUND (Dilution: 1:500, Santa Cruz, ref. sc-271938) or Histone H3 (Dilution: 1:1000, Cell Signaling, ref. 4620) were used as indicated.

#### RNA extraction & RNA-seq

After mechanical dissociation with Mixer Mill for 2 min at 30 Hz in 500 µL of Qiazol, RNA was extracted following Qiazol/chloroform extraction protocol (Qiagen, ref. 139046, MaXtract High density) and RNeasy Mini column (Qiagen, ref. 74106), following manufacturer instructions. RNA quality was checked on the Agilent TapeStation using RNA reagents. RNA extracts were sequenced on a NovaSeq 6000 (Illumina) in PE100 mode.

#### CITE-seq

Cells (1 × 10^6^) were blocked for 10 min at 4°C with 5 µL of human Trustain FcX (Biolegend, ref. 422301) and stained for 30 min at 4°C with 1 µg of TotalSeq A antibodies (Biolegend), using 1 barcoded antibody per sample for the MDAMB468 GOF VPR control, VPR FOSL1 gRNA2, VPR FOSL1 gRNA3, the HA control, and HA FOSL1. After washing, cells were filtered and counted. A pool of 5 stained samples was prepared by mixing 25,000 cells per sample (finale concentration of the pool, 2000 cells/µL). Approximately 16,000 cells were then loaded on a Chromium Single Cell Controller Instrument (Chromium Single Cell 3ʹv3, 10X Genomics, ref. PN-1000075) according to the manufacturer’s instructions. Samples and libraries were prepared according to the manufacturer’s instructions. Libraries were sequenced on a NovaSeq 6000 (Illumina) in PE 28-8-91 with a coverage of 50,000 reads/cell.

#### scRNA-seq

For each single-cell suspension or PDX dissociated cell sample^9^, approximately 3,000 cells were loaded on a Chromium Single Cell Controller Instrument according to the manufacturer’s instructions. Samples and libraries were prepared according to the manufacturer’s instructions. Libraries were sequenced on a NovaSeq 6000 (Illumina) in PE 28-8-91 with a coverage of 50,000 reads/cell.

#### CUT&Tag

The CUT&Tag-IT™ Assay Kit (Active Motif, ref. 53160) was used with 1:50 Anti-Rabbit antibodies for FOSL1 (Cell Signaling, ref. #5281), HA (Epicypher, ref. 13-2010) and H3K27ac (Cell Signaling, ref. #8173) following manufacturer instructions. The starting cell number was 500,000 cells for control cells and GOF cells, and down to 100,000 for persister cells after 35 days of 5 uM 5-FU treatment. The same digitonin concentration (0.05%) was maintained in all buffers, completing the kit with additional digitonin (Cell Signaling, ref. 16359). Sequencing libraries were prepared using Nextera™-Compatible Multiplex Primers (Active Motif, ref. 53155) and profiles were checked on the Agilent TapeStation using High-sensitivity D1000 reagents. CUT&Tag libraries were sequenced on a NovaSeq 6000 (Illumina) in PE100 mode.

#### Immunofluorescence

After media aspiration, cells in 24-well plates were washed with PBS and fixed with 3.7% formaldehyde in PBS for 15 min at room temperature. Cells were then washed with PBS three times, for 3 min each, and then permeabilized for 5 min at room temperature with 0.2% Triton X-100 in PBS. Samples were then incubated for 1 h at room temperature with a blocking buffer (1% SVF and 1% BSA in PBS) and then for 2 h at room temperature with 50 µL primary antibody anti-FOSL1 (dilution 1:100; Santa Cruz, ref. #sc-376148) in blocking buffer. After incubation, cells were washed three times with PBS with 0.2% Tween and then incubated for 1 h at room temperature in 50 µL secondary antibody solution consisting of the appropriate species-specific Alexa Fluor-conjugated antibodies (ThermoFisher, refs. A-21127, A-11008), diluted 1:1000 in blocking buffer. Coverslips were then washed three times with PBS with 0.2% Tween and mounted on slides with 10 µL ProLong Gold Antifade Reagent with DAPI (ThermoFisher, ref. P36971) at room temperature for 24 h. Confocal laser scanning microscopy (Carl Zeiss, LSM780) was used to scan GOF samples. Image processing was performed using Fiji Software, version 1.0. FOSL1 positive cells were evaluated using the “Analyze particles” option of Fiji Software using the following parameters: total cells (threshold=220 and pixel>50) and FOSL1+ cells (threshold=550 and pixel>50) for untreated cells and total cells (threshold=220 and pixel>100) and FOSL1+ cells (threshold=550 and pixel>100) for 5-FU treated cells.

#### Sample preparation for proteomics and mass spectrometry analysis

Lysis and wash buffers were prepared based on Hanson et al. 2016 protocol^72^. FOSL1 (Santa Cruz, ref. #sc28310 (C12)) and IgG (Sigma, ref. 12-371) immunoprecipitations on 1mg of lysates using protein A/G magnetic beads (Pierce, ref. 88802) were performed once in each cell condition after 7 days of DMSO or 5-FU. Beads were washed trice with 100 μL of 100 mM NH4HCO3 and resuspended in 100 μL of 100 mM NH4HCO3 before digestion by adding 0.2 μg of trypsin-LysC (Promega, ref. V5071) for 1 h at 37 °C. Samples were then loaded into custom-made C18 StageTips packed by stacking three AttractSPE disk (Affinisep, ref. SPE-Disks-Bio-C18-100.47.20) into a 200 µL micropipette tip for desalting. Peptides were eluted using a ratio of 40:60 CH3CN:H2O + 0.1% formic acid and vacuum concentrated to dryness with a SpeedVac device. Peptides were reconstituted in 10µl of injection buffer in 0.3% trifluoroacetic acid (TFA) before liquid chromatography-tandem mass spectrometry (LC-MS/MS) analysis.

#### LC-MS/MS analysis

Liquid chromatography (LC) was performed with a Vanquish Neo LC system (Thermo Scientific) coupled to an Orbitrap Astral mass spectrometer (MS), interfaced by a Nanospray Flex ion source (Thermo Scientific). Peptides were injected onto a C18 column (Thermo Scientific, ref. 75 µm inner diameter x 50 cm double nanoViper PepMap Neo, 2 μm, 100 Å) regulated at a temperature of 50°C, and separated with a linear gradient from 100% buffer A (100% H2O + 0,1% formic acid) to 28% buffer B (100% CH3CN + 0,1% formic acid) at a flow rate of 300 nL/min over 104 min. Peptides were analyzed in the MS applying a 2200V spray voltage, funnel RF level at 40% and a heated capillary temperature set to 285°C. MS full scans were recorded on the Orbitrap mass analyzer in centroid mode for ranges 380-980 m/z with a resolution of 240,000 at m/z 200, a normalized AGC target set at 500% and a maximum injection time of 5 ms. The MS2 acquisitions in the Astral analyzer were performed on a 150-2000 m/z range after fragmentation using HCD with 25% normalized collision energy and a normalized AGC target of 500%. A window width of 2 Da without overlap, 299 scan events and a maximum injection time of 3 ms were chosen for DIA. The loop control was set to 2.1 sec.

#### Persister cell counting

TNBC cells were plated in 24 multi-well plates at a density of 100,000 cells per well and treated with 5-FU for 7 days for MDAMB468 cells. Cultures were incubated in humidified 37°C incubators with an atmosphere of 5% CO_2_ in air, and the treated plates were monitored for growth using a microscope. Persister cells were evaluated after staining with Crystal Violet (Sigma, ref. C3886). In parallel, MDAMB468 persister cells were counted using a Countess automated cell counter (Invitrogen, ref. C10228) at the indicated time of treatment.

### Computational approaches

All statistical analyses were performed in R (v4.4) using custom R scripts. Code related to the following sections are deposited on Github (https://github.com/vallotlab/AP1_Multi_Chemopersistence_TNBC), and datasets are listed in **Table 1**.

#### Proteomics data processing

For identification, the data were searched against the Homo sapiens (UP000005640) Uniprot database using Pulsar search engine through Spectronaut v19 (Biognosys) by directDIA+ analysis using default search settings. Enzyme specificity was set to trypsin and a maximum of two missed cleavage sites was allowed. Carbamidomethyl cysteine, N-terminal acetylation and oxidation of methionine were set as variable modifications. The resulting files were further processed using myProMS v3.10. https://github.com/bioinfo-pf-curie/myproms^73^. For protein quantification, ion XICs from proteotypic peptides shared between compared conditions (TopN matching) were used, with missed cleavages and carbamidomethylation allowed. Median and scale normalization at peptide level was applied on the total signal to correct the XICs for each biological replicate (N = 5). To evaluate the statistical significance of the change in protein abundance, a linear model (adjusted on peptides and biological replicates) was performed, and a two sided T-test was apply on the fold change estimated by the model. The p-values were then adjusted using the Benjamini–Hochberg FDR procedure. In order to eliminate non-specific isolated proteins, only proteins were selected with at least three distinct peptides in three replicates out of five replicates, two-fold enrichment and an adjusted p-value ≤ 0.05 and for the unique proteins three distinct peptides. Proteins selected with these criteria were further used for quantitative data analysis.

#### Single-cell (sc) RNA-seq analysis

The scRNA-seq sequencing files were preprocessed using the cellRanger pipeline. For PDX samples, files were aligned to the hg38 and mm10 genomes, and only cells with a majority of human reads (> 75%) were retained for the analysis. For the MDAMB468, HCC38, and BT20 human cell lines, sequences were aligned to the hg38 genome. For all datasets, cells were kept within a 100,00 read coverage limit. For PDX samples, cells had a coverage between 2,000 and 9,000 genes, with mitochondrial reads below 25%. For MDAMB468, HCC38, and BT20 cells, cells had a coverage between 3,000 genes and 8,000 genes, with mitochondrial reads below 15%. Normalization (SCTransform method), dimensionality reduction, and Louvain clustering were performed using Seurat (v5.1.0), keeping the first 30 principal components (PC). The cell cycle was scored for each cell using Seurat. For the PDXs, BT20, HCC38, and MDAMB468 models, persister cells were compared to cells from the untreated model. For each model, differentially expressed genes were computed using the “FindMarkers” Seurat function with default parameters. UMAP plots of gene expression were obtained using Nebulosa (v1.14.0) to recover the signal from dropped-out features using kernel density estimation.

#### Bulk RNA-seq analysis and microarray analysis

The bulk RNA-seq samples were preprocessed using the Nextflow raw-qc analysis pipeline from Institut Curie (<u>10.5281/zenodo.7515639</u>). PDX samples were aligned on hg38 and mm10 genomes, and only reads aligning to the hg38 genome were retained for the analysis. Normalization was made using the TMM method from the edgeR package (v4.2.0)^74,75^. U133 Affymetrix microarrays were preprocessed with the GCRMA package^76^.

#### Integrated analysis of microarray, bulk RNA-seq, and scRNA-seq samples

Raw count matrices of scRNA-seq samples were first merged, keeping all features (and putting 0 for missing values). A pseudo-bulk count matrix was then created by aggregating all cell counts coming from the same tumor sample with Presto (v1.0.0). After this step, all raw count matrices coming from microarray, bulk RNA-seq, and scRNA-seq samples were merged together, whereby a set of features common to all samples was maintained. LogNormalisation was performed using the TMM method from the edgeR package (v4.2.0). Sample and technology effects were removed using the removeBatcheffect function from Limma (3.60.3)^28^, which gave a corrected logcount matrix. Principal component analysis (PCA) was applied on the 500 most variable features within this matrix. For the differential analysis, the voom method from limma was used on the raw matrix count, taking into account in the fitted linear model the group of the sample (persister, chemo-naïve, or relapse) and the model (PDX of origin). The persister and chemo-naïve groups were compared using the makeContrasts function.

#### Regulatory network inference

Regulon activity was inferred using DecoupleR (v2.10.0)^32^. The network of transcription factors (TFs) and their transcriptional targets were retrieved from the CollecTRI database using OmnipathR (v3.11.10), with the option “split_complexes = False”. With this network, a consensus score of TF enrichment scores was obtained by combining 3 methods (ulm, wsum and mlm) on the T-value from the differential expression analysis obtained with limma. T-values were either directly retrieved from the differential expression analysis when limma was used, or calculated by -log10(p.adj)*log2FC when Seurat was used. The TFs with the top 10 scores were selected. Regulon activity per sample per TF/complex was computed using the normalized weighted sum method on the corrected logcount matrix obtained with limma.

#### CUT&Tag analysis

Raw sequencing files were processed using an in-house pipeline (https://github.com/vallotlab/bulk_Epigenomics). Raw reads were mapped in paired end mode using BowTie2 with options "-k 1 -m 1" on the human genome (hg38). Peak calling by condition and antibody used (histone modification or transcription factor binding) was performed on bam files using macs3^77^, with the following parameters: "--max-gap 500 -f BAM -q 0.000001 --nolambda". PCA and differential analysis were performed from count matrices on a merged peak set from the different conditions (untreated, 5-FU, or FOSL1 overexpression experiments) and antibodies (anti-H3K27ac, anti-FOSL1, or anti-HA). Peaks covered with less than 5 reads across all samples were filtered out. Normalization and differential analyses were performed with DESeq2 (v1.44.0)^78^. PCA were calculated on the log-normalized count matrices selected for the 500 most variable peaks. To compare H3K27ac and FOSL1 enrichments, a common set of consensus peaks was assembled by combining peaks from all conditions (chemo-naïve, 5-FU, or GOF cells) for both antibodies. Profile plots for CUT&Tag were made using Plotgardener (v1.10.2)^79^. For each snapshot, consensus peaks, associated log2 fold-change (log2FC) and adj. *P* computed with DESeq2s are indicated on the plots.

#### Motif enrichment analysis

Motif enrichment analysis was performed on differentially enriched peaks for H3K27ac, comparing 5-FU–treated and wild-type MDAMB468 cells. The Hocomoco database (v12) and Memes (v1.12.0) were used to perform motif enrichment on the top 1000 differentially enriched peaks (ordered by decreasing T-values). A set of 5000 random peaks with no enrichment upon 5-FU treatment was used as background.

## DATA AVAILABILITY

The datasets described in this study are deposited in GEO repository GSE280454 and GSE280455. The mass spectrometry proteomics raw data will be deposited to the ProteomeXchange Consortium via the PRIDE partner repository^80^.

## Authors’ contributions

LB, JM performed experiments. PDX experiments were performed by EM, LS, AD and EM. CH and JM performed lentivirus production. GJ, PP, MS, SG and CV performed omics data analysis. FD carried out the MS experimental work and DL supervised MS and data analysis. CV and JM conceived and designed experiments. CV, JM, LB and GJ wrote the manuscript with input from all authors.

## Acknowledgments

pHIV-SFFV-MCS-IRES-GFP was kindly provided by Jérémy Welsch and Caroline COSTA from AniRA Vectorologie in Lyon, France. Lentivirus particles for Cas9WT-P2A-mCherry (Addgene-ref. 99154) and dCas9-VPR-P2A-mCherry (Addgene, ref. 154193) were provided by the CRISPR’IT platform of Institut Curie. Next-Generation Sequencing platform of Institut Curie performed sequencing. Microarray preprocessing was performed by Pierre Gestraud at the U900 Bioinformatics Platform of Institut Curie. The results on human tumor samples shown here are based upon data generated by the TCGA Research Network: https://www.cancer.gov/tcga. The LSMP thanks Patrick Poullet from the bioinformatics platform of the Institut Curie U900 for the continuous development of myProMS.

## Additional Information

Correspondence and requests for materials should be addressed to.

